# Value Shapes Abstraction During Learning

**DOI:** 10.1101/2020.10.29.361469

**Authors:** Aurelio Cortese, Asuka Yamamoto, Maryam Hashemzadeh, Pradyumna Sepulveda, Mitsuo Kawato, Benedetto De Martino

## Abstract

The human brain excels at constructing and using abstractions, such as rules, or concepts. Here, in two fMRI experiments, we demonstrate a mechanism of abstraction built upon the valuation of sensory features. Human volunteers learned novel association rules linking simple visual features. Mixture-of-experts reinforcement learning algorithms revealed that, with learning, high-value abstract representations increasingly guided participants’ behaviour, resulting in better choices and higher subjective confidence. We also found that the brain area computing value signals - the ventromedial prefrontal cortex – prioritized and selected latent task elements during abstraction, both locally and through its connection to the visual cortex. Such coding scheme predicts a causal role for valuation: in a second experiment, we used multivoxel neural reinforcement to test for the causality of feature valuation in the sensory cortex as a mechanism of abstraction. Tagging the neural representation of a task’s feature with rewards evoked abstraction-based decisions. Together, these findings provide a new interpretation of value as a goal-dependent, key factor in forging abstract representations.

## Introduction

### “All art is an abstraction to some degree.” Henry Moore

Art is one of the best examples of abstraction, the unique ability of the human mind to organise information beyond the immediate sensory reality. Abstraction is by no means restricted to high level cognitive behaviour such as art production: it envelops every aspect of our interaction with the environment. Imagine that you are hiking in a park, and you need to cross a stream. Albeit deceptively simple, this scenario already requires the processing of a myriad visual (and auditory, etc.) features. For an agent that operates directly on each feature in this complex sensory space, any meaningful behavioural trajectory (such as crossing a stream) would quickly involve intractable computations. This is well exemplified in reinforcement learning (RL), where in complex and/or multidimensional problems, classic RL algorithms rapidly collapse (1–3). If, on the other hand, the agent is able to first ‘abstract’ the current state to a lower dimensional manifold, representing only relevant features, behaviour would be far more flexible and efficient (4–6). Attention (7–9), and more generally, the ability to act upon subspaces, concepts or abstract representations have been proposed as effective solutions to overcome the computational bottlenecks arising from sensory level operations in RL (4, 5, 10–12). Abstractions can be thus thought of as simplified maps carved from higher dimensional space in which details have been removed or transformed, in order to focus on a subset of interconnected features, i.e., a higher-order concept, category or schema (13, 14).

How are abstract representations constructed in the human brain? For flexible deployment, abstraction should depend on task goals. From a psychological or neuroeconomic point of view, task goals generally determine what is valuable (15–17), such that if I need to light a fire, matches are much more valuable than a glass of water. Hence, we hypothesized that valuation processes are directly related to abstraction.

Value representations have been linked to neural activity in the ventromedial prefrontal cortex (vmPFC) in the context of economic choices (16, 18). More recently, vmPFC role has also been extended to the computation of confidence (19–22). While this line of work has been extremely fruitful, it has mostly focused on the hedonic and rewarding aspect of value instead of its broader functional role. In the field of memory, a large corpus of work has shown that vmPFC is crucial for the formation of schemas or conceptual knowledge (13, 14, 23–25), as well as generalizations (26). The vmPFC also collates goal-relevant information from elsewhere in the brain (32). Considering its connectivity pattern (27), the vmPFC is well suited to play a pivotal role in the circuit that involves the hippocampal formation (HPC) and orbitofrontal cortex (OFC), dedicated to extracting latent task information and regularities important for navigating behavioural goals (6, 28–31). Thus, the aim of this study is twofold: i) demonstrate abstraction emerges during the course of learning, and ii) investigate how the brain, and specifically the vmPFC, uses valuation upon low-level sensory features to forge abstract representations.

To achieve this, we leveraged a task in which human participants repeatedly learned novel association rules, while their brain activity was recorded with fMRI. Reinforcement learning (RL) and mixture-of-experts modelling (33, 34) allowed us to track participants’ valuation processes and to dissociate their learning strategies (both at the behavioural and neural levels) based on the degree of abstraction. Participants reported confidence in their performance positively correlated with their ability to abstract. In a second experiment, we studied the causal role of value in promoting abstraction through directed effect in sensory cortices. To anticipate our results, we show that vmPFC and its connection to visual cortex construct abstract representations through a goal-dependent valuation process, that is implemented as top-down control of sensory cortices.

## Results

### Experimental design

The goal of the learning task was to present a problem that could be solved according to two strategies, based on the sampled task-space dimensionality. A simple, slower strategy akin to pattern recognition, and a more sophisticated one that required abstraction to use the underlying structure. Participants (N = 33) learned the fruit preference of pacman-like characters formed by the combination of 3 visual features (colour, mouth direction and stripes orientation, Figure 1A-B). The preference was governed by the combination of two features, selected randomly by our computer program on each block (Figure 1A-B). Learning the blocks’ rules essentially required to uncover the hidden associations between features and fruits. Although participants were instructed that one feature was irrelevant, they did not know which. A block ended either when a sequence of 8-12 (randomly set by our computer program) correct choices was detected or upon reaching its upper limit (80 trials). Variable stopping criteria were used to avoid participants learning that a fixed sequence was predictive of block ending. On each trial, participants could see the outcome after selecting a fruit. A green box appeared around the chosen fruit if the choice was correct (red otherwise). Additionally, participants were instructed that the faster they learned a block’s rule, the higher the reward. At the end of the session, a final reward was delivered, commensurate to one’s performance (see Methods).

**Figure 1:**
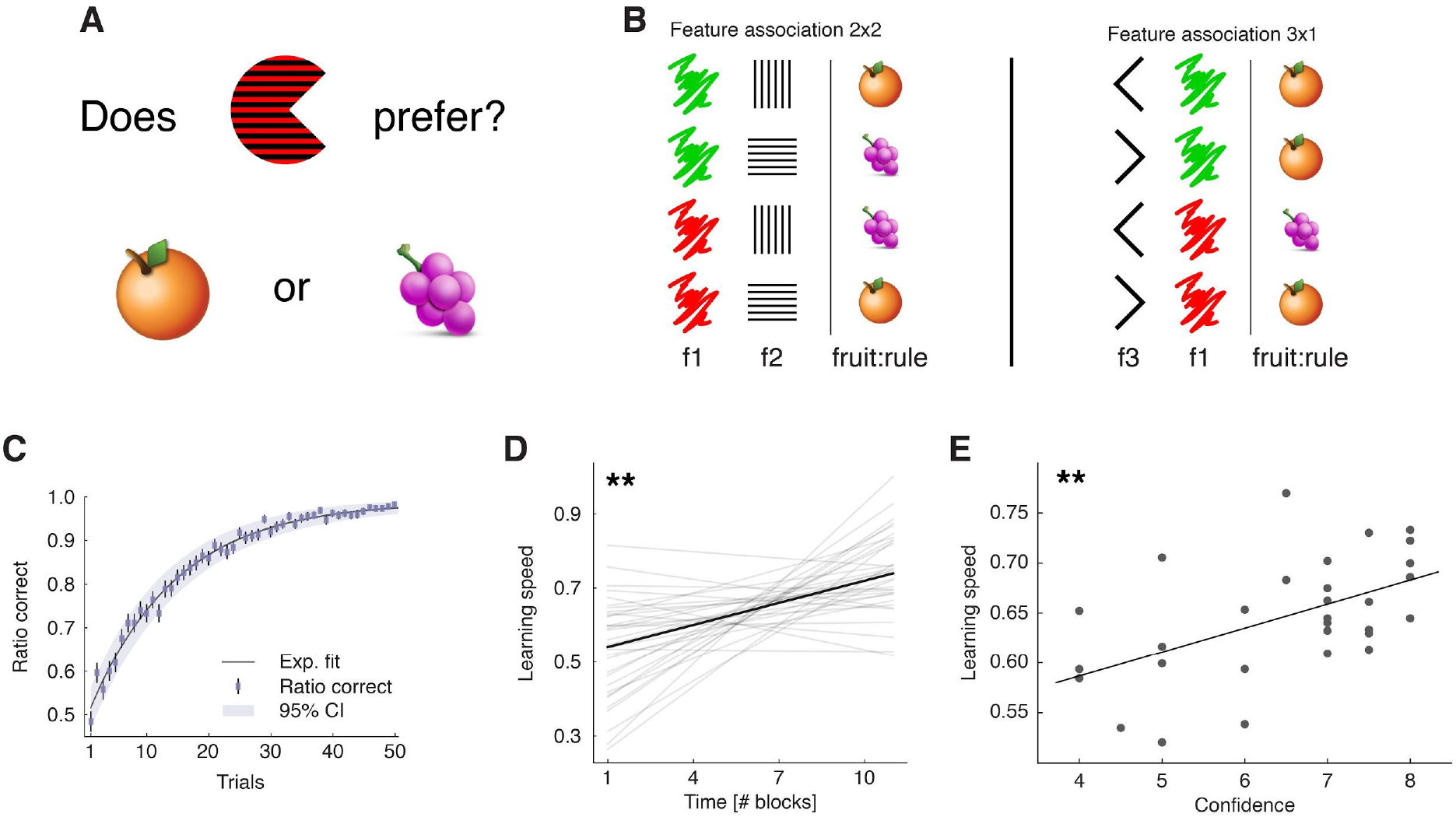
Learning task and behavioural results. ***A***, Task: participants learned the fruit preference of pacman-like characters, which changed on each block. ***B***, Examples of the two types of fruit associations. The 4 combinations arising from 2 features with 2 levels were divided into symmetric (2×2) and asymmetric (3×1) cases. f1-3: features 1 to 3; fruit:rule refers to the fruit as being the association rule. The two block types were included to avoid participants learning the rules by simple deduction. If all blocks had symmetric association rules and participants knew this, they could simply learn one feature-fruit association (e.g., green-vertical), and from there deduce all other combinations. ***C***, Trial-by-trial ratio correct as a measure of within-block learning. Dots represent the mean across participants, while the error bars the SEM, and the shaded area the 95% CI (N = 33). Participant-level ratio correct was computed for each trial across all completed blocks. ***D***, Correlation between learning speed and time, across participants. The learning speed was computed as the inverse of the max-normalized number of trials taken to complete a block. The thin grey lines represent individual participants least square fits, while the black line represents the group-average fit. The correlation was computed with group-averaged data points (N = 11). ***E***, correlation between confidence judgements and learning speed, across participants. Each dot represents data from one participant, and the thick line the regression fit (N = 31 [2 missing data]), ** *p*<0.01

### Behavioural accounts of learning

We verified that participants learned the task sensibly. Within blocks, performance was higher than chance already from the second trial (Figure 1C, one-sample t-test against mean of 0.5, trial 2: *t*_32_ = 4.13, *P*_*(FDR)*_ < 10^−3^, trial 3: *t*_32_ = 2.47, *P*_*(FDR)*_ = 0.014, all trials t>3: *P*_*(FDR)*_ < 10^−3^). Considering the whole experimental session, learning speed (i.e., how quickly participants completed a given block) significantly increased across blocks (Figure 1D, N = 11 time points, Pearson’s *r* = 0.80, *p* = 0.003). These results not only confirmed participants learned the task rule in each block, but also that they learned to use the more efficient strategy. Of note, in this task, the only way to solve the blocks faster is by using the correct subset of dimensions (the abstract representation). When asked at the end of the session their degree of confidence in having performed the task well, participants’ self-reports correlated with their learning speed (N = 31 [2 missing data], robust regression slope = 0.024, *t*_29_ = 3.27, *p* = 0.003, Figure 1E), but not with the overall number of trials, or the product of the proportion of successes (learning speed: Pearson’s *r* = 0.49, *p* = 0.024, total trials: *r* = −0.13, *p* = 0.47, test for difference in *r*: z = 2.49, *p* = 0.013; product of the proportion of successes: *r* = −0.06, *p* = 0.75, test for difference in *r*: z = 2.23, *p* = 0.026). We also confirmed that the block type (defined by the relevant features, e.g., colour-orientation) or the association type (e.g., symmetric 2×2) did not systematically affect the learning speed (Figure S1).

### Mixture-of-experts reinforcement learning for the discovery of abstract representations

Was participants’ learning behaviour guided by the selection of accurate representations? To this end, we built upon a classic RL algorithm (Q-learning) (35) in which state-action value functions (beliefs), used to make predictions on future rewards, are updated according to a trial’s task state and action’s outcome. In this study, task states were defined by the number of features combinations that the agent may track; hence, we devised algorithms that differed in their state-space dimensionality. The first algorithm, called Feature RL, explicitly tracks all combinations of 3 features, 2^3^ = 8 states (Figure 2A, top left). This algorithm is anchored at a low feature level because each combination of the 3 features results in a unique fingerprint - one simply learns direct pairings between visual patterns and fruits (actions). Conversely, a second algorithm, called Abstract RL, uses a more compact or *abstract state representation* in which only two features are tracked. These compressed representations reduce the explored state-space by half, 2^2^ = 4 states (Figure 2A, top right). Importantly, in this task environment there can be as many as 3 Abstract RL in parallel, one for each combination of two features.

**Figure 2:**
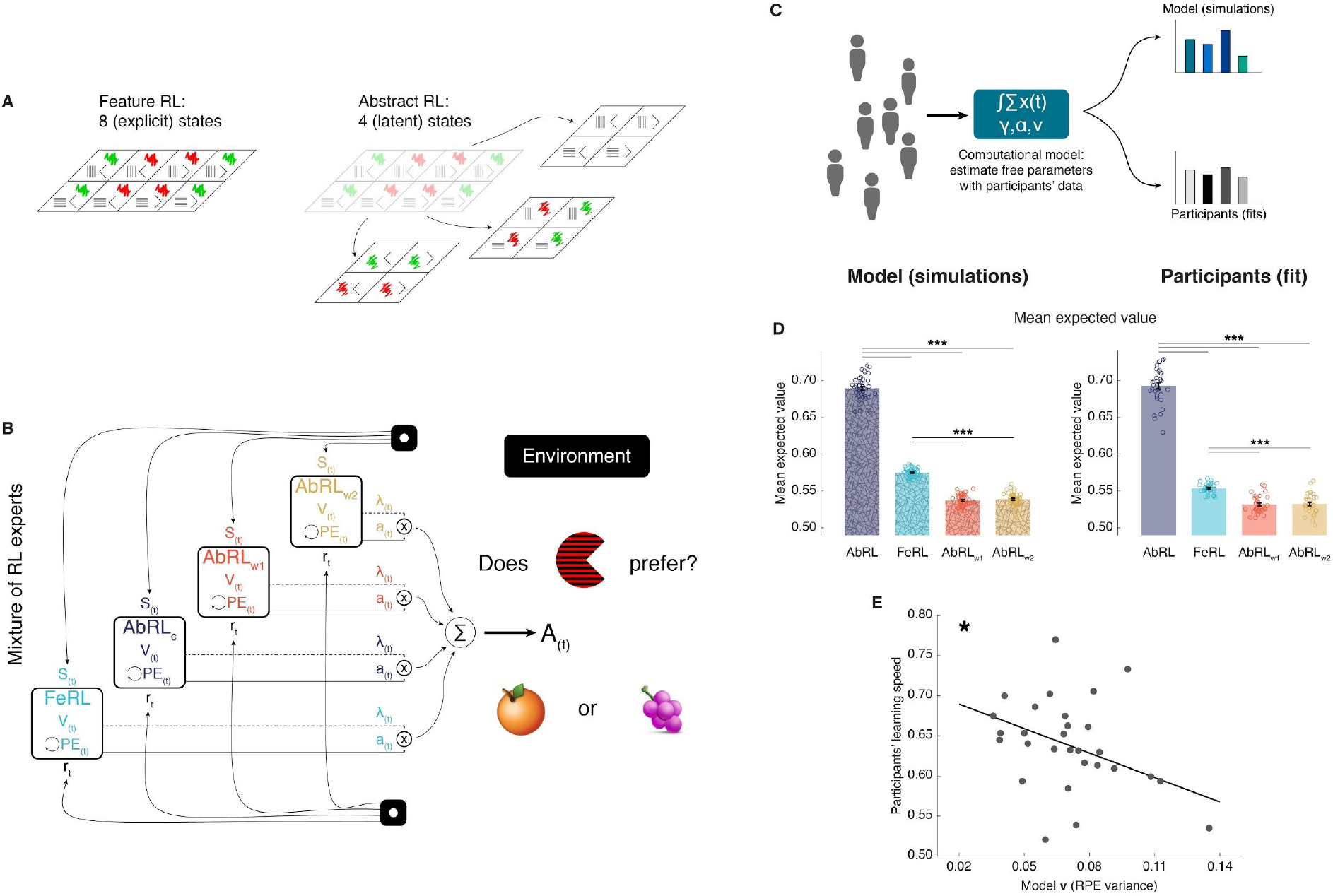
Mixture of reinforcement learning (RL) experts and value computation. ***A***, Outline of the representational spaces of each RL algorithm comprising the mixture-of-experts architecture. ***B***, Illustration of the model architecture. See Methods for a formal description of the model. All experts had the same number of hyperparameters: the learning rate © (how much the latest outcome affect the agent’s beliefs), the forgetting factor © (how much prior RPEs influence current decisions), and the RPE variance *v*, modulating the sharpness with which the mixture-of-expert RL model should favour the best performing algorithm in the current trial. ***C***, Approach used for data analysis and model simulation. The model was first fit to participants’ data with Hierarchical Bayesian Inference (39). Estimated hyperparameters were used to compute the value functions of participants’ data, as well as to generate new, artificial choice data and compute simulated value functions. ***D***, Averaged expected value across all states for the chosen action in each RL expert. Left: simulated data, right: participants’ empirical data. Dots represent individual agents (left) or participants (right), bars the mean and error bars the SEM. Statistical comparison was performed with two-sided Wilcoxon signed rank tests. Model: AbRL_w1_ vs AbRL_w2_, z = 1.00, *p* = 0.32, all remaining comparisons, z = 5.84, \*\*\**p*<0.001; Participants: AbRL_w1_ vs AbRL_w2_, z = −0.08, *p* = 0.94, all remaining comparisons, z ≥ 4.92, \*\*\**p*<0.001. AbRL: Abstract RL, FeRL: Feature RL, AbRL_w1_: wrong-1 Abstract RL, AbRL_w2_: wrong-2 Abstract RL. ***E***, Correlation between RPE variance and learning speed (outliers removed, N = 29). Dots represent individual participants’ data, the thick line the regression fit, * *p*<0.05.

The above four RL algorithms were combined in a mixture-of-experts architecture (33, 34, 36), Figure 2B and Methods. The key intuition behind this approach is that, at the beginning of a new block, the agent does not know which abstract representation is correct (i.e., which features are relevant). Thus, the agent should learn which representations are most predictive of reward, and thereby exploit the best representation for action selection. While all experts participated in action selection, their learning uncertainty (RPE: reward prediction error) determined the strength in doing so (34, 37, 38). This architecture allowed us to track the value function of each RL expert separately, while using a unique, global action on each trial.

Estimated hyperparameters were used to compute the value functions of participants’ data, as well as to generate new, artificial choice data and value functions (Figure 2C, and Methods). Simulations indicated expected value was highest for the appropriate Abstract RL, followed by Feature RL, and the two Abstract RLs based on irrelevant features as the lowest (Figure 2D). Participants’ empirical data displayed the same pattern, whereby the value function of the appropriate Abstract RL was higher than other RL algorithms (Figure 2D, right side). Note the large difference between appropriate Abstract RL and Feature RL: this is due to the appropriate Abstract RL being an ‘oracle’, as it has access to the correct low-dimensional state from the beginning. The RPE variance ***v*** adjusted the sharpness with which each RL’s (un)certainty was considered for the expert weighting. Crucially, the variance ***v*** was associated with participants’ learning speed, such that participants who learned the blocks’ rules quickly were sharper in expert selection (Figure 2E, N = 29, robust regression slope = − 1.02, *t*_27_ = −2.59, *p* = 0.015). These modelling results provided a first layer of computational support for the hypothesis that valuation is related to abstractions.

### Behaviour shifts from Feature- to Abstract-based reinforcement learning

The mixture-of-expert RL model uncovered that participants who learned faster relied more on the best RL model value representations. Critically, the modelling established that choices were mostly driven by either the appropriate Abstract RL or Feature RL - as they had higher expected values (although note that the other Abstract RLs had mean values greater than a null level of 0.5). Hence, we next explicitly explained participants’ choices and learning according to either strategy. Here, given the task space (Figure 2A), the only way to solve a block’s rule faster is to use abstract representations. As such, we expect to observe a shift from Feature RL towards Abstract RL with learning.

Both algorithms had two hyperparameters: the learning rate ***α*** and greediness ***β*** (inverse temperature - the strength that expected value has on determining actions). Using the estimated hyperparameters, we generated new, synthetic data to evaluate how fast artificial agents, implementing either Feature RL or Abstract RL, solved the learning task (see Methods). The simulations attested Feature RL was slower and less efficient: yielding lower learning speed (Figure 3A left, two-sided Wilcoxon rank sum test, z = 90.97, *p* < 10^−30^), and a higher percentage of failed blocks (Figure 3A right, two-sided Wilcoxon rank sum test, z = 8.16, *p* < 10^−6^).

**Figure 3:**
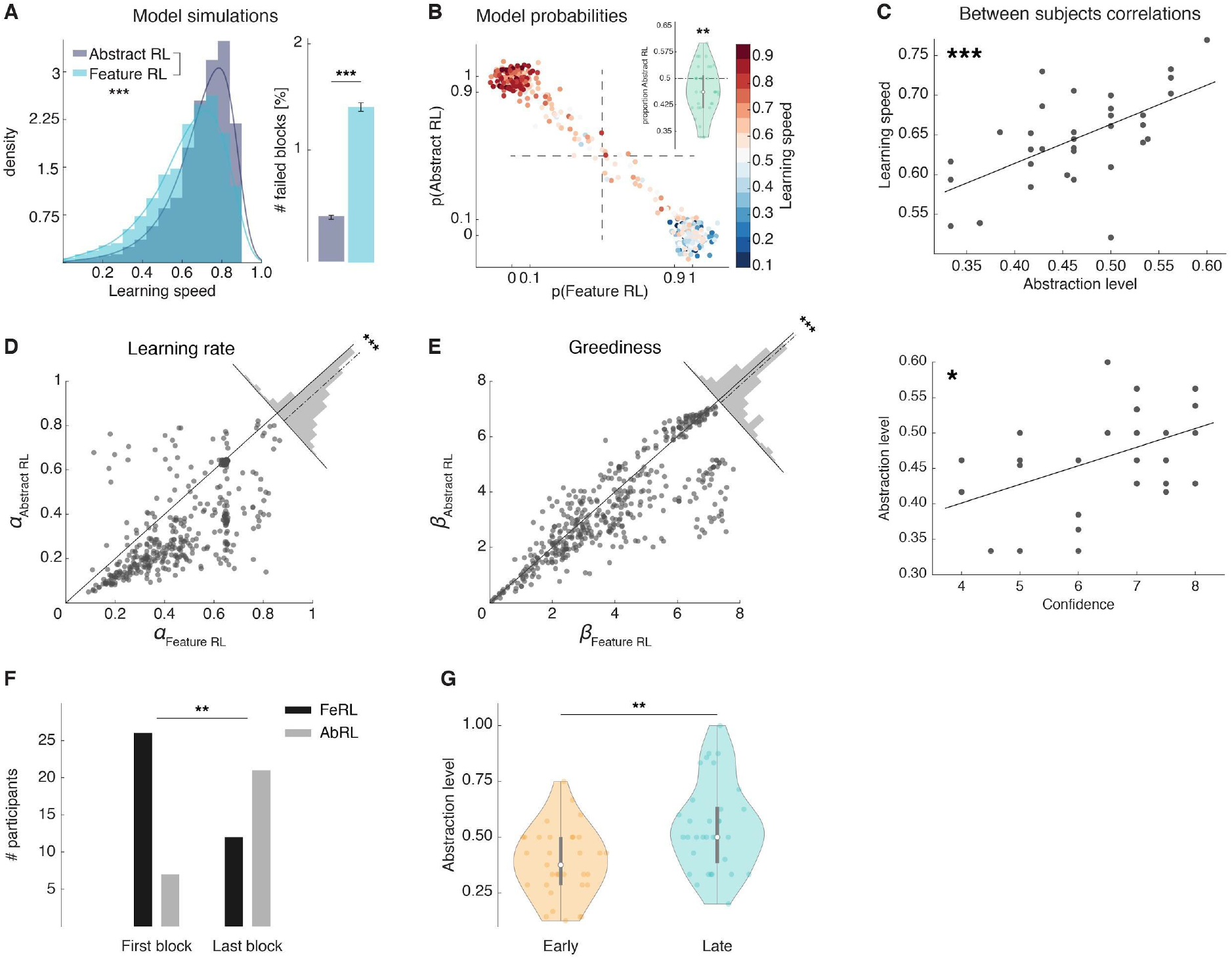
Feature RL vs Abstract RL are related to learning speed. ***A***, Simulated learning speed and % of failed blocks for both Abstract RL and Feature RL. To make the simulations more realistic, arbitrary noise was injected into the simulation, altering the state (see Methods). N = 500 simulations of 45 agents. Right plot: bars represent the mean, error bars the SEM. ***B***, Relationship between block-by-block best-fitting model and learning speed in participants. Each dot represents one block, from one participant. Data aggregated across all participants. Note some dots fall beyond p=1 or p=0: the effect is due to dots being scattered with noise in their x-y coordinates for better visualization. ***C***, Between participants correlations. Top: abstraction level vs learning speed. The abstraction level was computed as the average over all blocks completed by a given participant (code: Feature RL = 0, Abstract RL = 1). Bottom: confidence vs abstraction level. Dots represent individual participants (top: N = 33, bottom: N = 31, some dots are overlapping). ***D***, Learning rate was not symmetrically distributed across the two algorithms. ***E***, Greediness was not symmetrically distributed across the two algorithms. For both ***D*** and ***E***, each dot represents one block, from one participant, data aggregated across all participants. Histograms represent the distribution of the data around the midline. ***F***, Number of participants for which Feature RL or Abstract RL explained best their choice behaviour in the first and last block of the experimental session. ***G***, Abstraction level was computed separately with blocks from the first half (early) and latter half (late) session. * *p*<0.05, ** *p*<0.01, *** *p*<0.001.

Model comparison at the single participant and block level (39) provided a direct way to infer which algorithm was more likely to explain learning in any given block. Overall, there were similar proportions of blocks classified as Feature RL and Abstract RL. This indicates participants used both learning strategies (binomial test applied to all blocks: proportion of Abstract RL = 0.47 vs. equal level = 0.5, *P*(212|449) = 0.26, Figure 3B; two-sided t-test of participant-level proportions: lower but close to 0.5, *t*_32_ = −2.87, *p* = 0.007, Figure 3B inset).

As suggested by the simulations (Figure 3A), the strategy that best explained participants’ block data accounted for the distribution of learning speed measures in each block. Where learning proceeded slowly, Feature RL was consistently predominant (Figure 3B), while the reverse happened in blocks where participants displayed fast learning (Figure 3B). Across participants, the degree of abstraction (propensity to use Abstract RL) correlated with the empirical learning speed (N = 33, robust regression, slope = 0.52, *t*_31_ = 4.56, *p* = 7.64×10^−5^, Figure 3C top). Participants’ reported sense of confidence in having performed the task well also significantly correlated with the degree of abstraction (N = 31, robust regression, slope = 0.026, *t*_29_ = 2.69, *p* = 0.012, Figure 3C, bottom). Together with confidence self-reports being predictive of learning speed (Figure 1E), these results raise intriguing questions on the function of metacognition, as participants appeared to grasp their own ability to construct and use abstractions (40).

The two RL algorithms revealed a second aspect of learning. Feature RL had consistently higher learning rates ***α*** compared with Abstract RL (two-sided Wilcoxon rank sum test against median 0, z = 14.33, *p* < 10^−30^, Figure 3D). This difference can be explained intuitively as follows. Integration of information has to happen over a longer time horizon in Abstract RL, because a single trial per se is uninformative with respect to the rule. Conversely, a higher learning rate would allow the agent to make larger updates on states that were less frequently visited, as would happen in Feature RL. A similar asymmetry was found with greediness (Figure 3E, two-sided Wilcoxon rank sum test against median 0, z = 7.14, *p* < 10^−10^), suggesting action selection tended to be more deterministic in Feature RL (i.e., large ***β***).

We predicted that abstraction use should increase with learning, because the brain has to construct abstractions in the first place and will thus initially rely on Feature RL. To test this hypothesis, we quantified the number of participants using Feature RL or Abstract RL strategy in their first and last blocks. On their first block, most participants relied on Feature RL, while the pattern reversed in the last block (two-sided sign test, z = −2.77, *p* = 0.006, Figure 3F). Computing the abstraction level separately for the session’s median split of early and late blocks also resulted in higher abstraction in the late blocks (two-sided sign test, z = −2.94, *p* = 0.003, Figure 3G). This coarse effect was complemented by a linear increase towards higher abstraction from early to late blocks (Figure S2A).

Supporting the current modelling framework, the mean expected value of the chosen action was higher for Abstract RL (Figure S2B), and model hyperparameters could be recovered in the presence of noise (Figure S3, see Methods) (41).

### The role of vmPFC in constructing goal-dependent value from sensory features

The computational approach confirmed participants relied on both a low-level, feature strategy, and a more sophisticated abstract strategy (i.e., Feature RL and Abstract RL; Figure 2D, 3B). Besides proving that abstract representations were generally associated with higher expected value, the modelling approach further allowed us to explicitly *classify* trials as belonging to either learning strategy. Here, our goal was to dissociate the neural signatures of these distinct learning strategies in order to elucidate how abstract representations are constructed by the human brain.

First, we reasoned that an anticipatory value signal might emerge in the vmPFC at stimulus presentation (42). We performed a general linear model (GLM) analysis of the neuroimaging data with regressors for ‘High value’ and ‘Low value’ trials, elected by the block-level best fitting algorithm (Feature RL or Abstract RL, while controlling for other confounding factors, see Methods for full GLM and regressors specification). As predicted, activity in the vmPFC strongly correlated with value magnitude (Figure 4A). That is, the vmPFC indexed the anticipated value constructed from the pacman features at stimulus presentation time. We used this signal to functionally define, for ensuing analyses, the subregion of the vmPFC that was maximally related to task computations about value when (pacman’s) visual features were integrated. Concurrently, activity in insular and dorsal prefrontal cortices increased under condition of low expected value. This pattern of activity is consistent with previous studies on error monitoring and processing (43, 44) (Figure S4).

**Figure 4:**
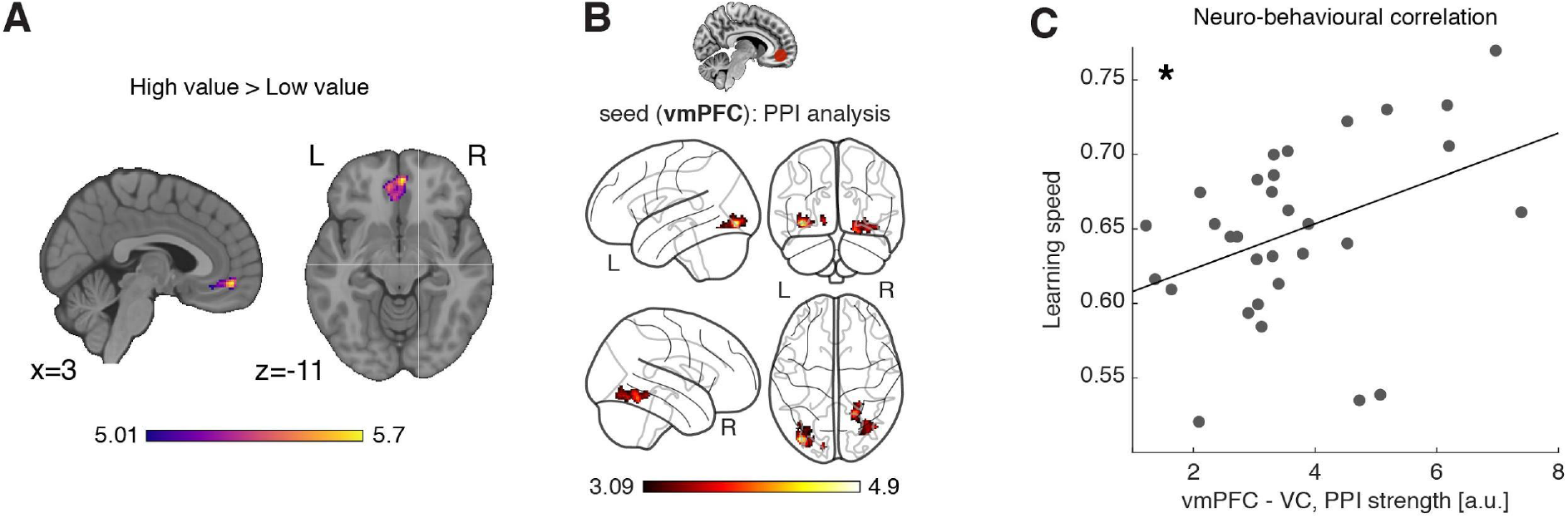
Neural substrates of value construction during learning. ***A***, Correlates of anticipated value at pacman stimulus presentation time. Trials were labelled according to a median split of the expected value for the chosen action as computed by the best fitting model - Feature RL or Abstract RL - at the block level. Mass univariate analysis, contrast ‘High value’ > ‘Low value’. vmPFC peak at [2 50 −10]. The statistical parametric map was z-transformed and plotted at *p*(FWE) < 0.05. ***B***, Psychophysiological interaction, using as seed a sphere (radius = 6mm) centred around the participant-specific peak voxel, constrained within a 25mm sphere centred around the group-level peak coordinate from contrast in (***A***). The statistical parametric map was z-transformed and plotted at *p*(fpr) < 0.001. ***C***, The strength of the interaction between the vmPFC and VC was predictive of the participants’ ability to learn the blocks’ rules. Dots represent individual subject’s data points, the line a regression fit, * *p*<0.05.

In order for the vmPFC to construct goal-dependent value signals, it should receive relevant feature information from other brain areas. Specifically, from visual cortices given the nature of our task. Thus, we computed a psychophysiological interaction (PPI) analysis (45), to isolate regions whose functional coupling with the vmPFC at the time of stimulus presentation was dependent on the magnitude of expected value. Supporting the idea that vmPFC based its predictions on the integration of visual features, only the connectivity between VC and vmPFC was higher on trials that carried large expected value compared to low value trials (Figure 4B). Strikingly, the strength of this VC - vmPFC interaction was predictive of the overall learning speed across participants (N = 31, robust regression, slope = 0.016, *t*_29_ = 2.53, *p* = 0.016, Figure 4C), such that participants who had a stronger modulation of the coupling between the vmPFC and VC also learned the blocks’ rules faster.

### Value-sensitive vmPFC subregion prioritizes abstract elements

Having established that the vmPFC computes a goal-dependent value signal, we evaluated whether this region’s activity level was sensitive to the strategy participants used. To do so, we used the same GLM introduced earlier, and estimated two new statistical maps from the regressors ‘Abstract RL’ and ‘Feature RL’ (see Methods and Supplementary note 1). We extracted the peak activity at the participant level, under Feature RL and Abstract RL conditions, in two regions-of-interest (ROI). Specifically, we focused on the vmPFC and the HPC, as both have been consistently linked with abstraction, feature-based and conceptual learning. The HPC was defined anatomically (Figure 5A top), while the vmPFC was defined as the voxels sensitive to the orthogonal contrast ‘High value’ > ‘Low value’ from the same GLM (Figure 5A bottom). A linear mixed effects model (LMEM) with fixed effects ‘ROI’ and ‘strategy’ [LMEM: ‘y ~ ROI * strategy + (1|participants)’, y: ROI activity] revealed significant main effects of ‘ROI’ (*t*_128_ = 2.16, *p* = 0.033), and ‘strategy’ (*t*_128_ = 3.07, *p* = 0.003), and a significant interaction (*t*_128_ = −2.29, *p* = 0.024), illustrating different HPC and vmPFC recruitment (Figure 5B). Post-hoc comparisons showed vmPFC activity levels distinguished well Feature RL and Abstract RL cases (LMEM: *t*_64_ = 2.94, *p*_*(FDR)*_ = 0.009), while the HPC remained agnostic to the style of learning (LMEM: *t*_64_ = 0.62, *p*_*(FDR)*_ = 0.54). Alternative explanations were unlikely, as there was no effect in terms of both the simple correlation between value-type trials and algorithms, and task difficulty measured by reaction times (Figure S5).

**Figure 5:**
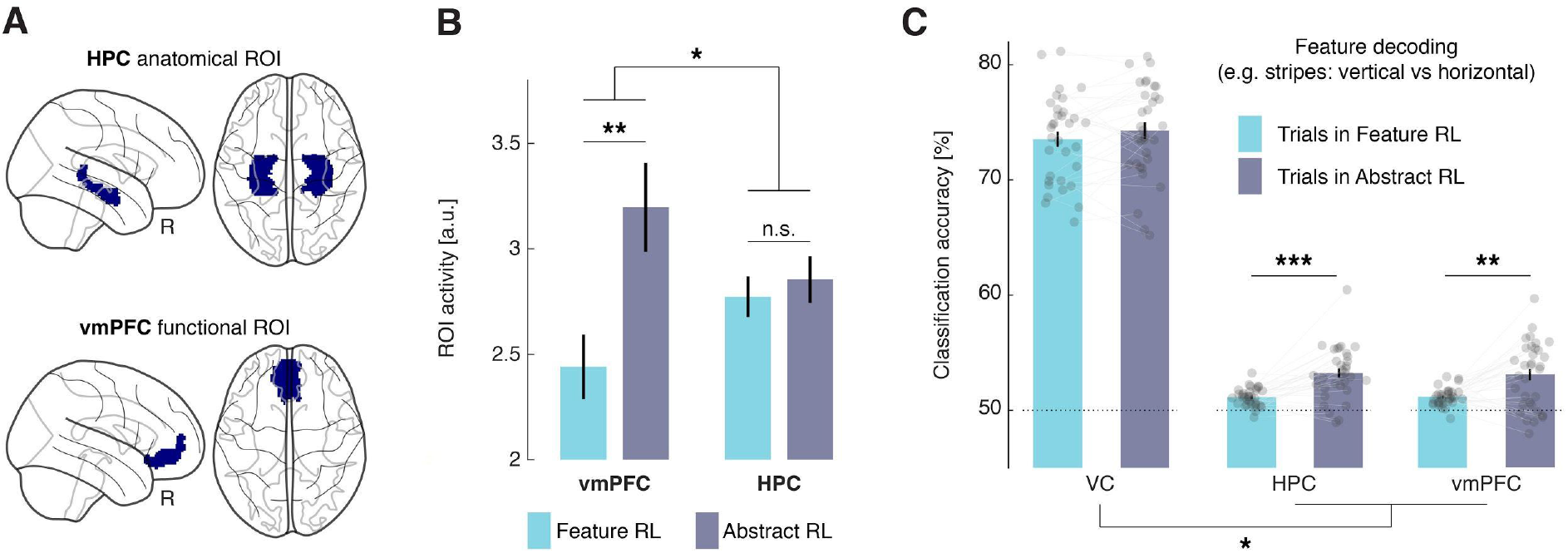
Neural substrate of abstraction. ***A***, Regions of interest for univariate and multivariate analyses. The HPC was defined through anatomical automated labelling (FreeSurfer). The vmPFC was functionally defined as the cluster of voxels found with the orthogonal contrast ‘High value’ > ‘Low value’, at *P*(unc) < 0.0001. ***B***, ROI activity levels corresponding to each learning mode were extracted from the contrasts ‘Feature RL’ > ‘Abstract RL’, and ‘Abstract RL’ > ‘Feature RL’. Coloured bars represent the mean, error bars the SEM. ***C***, Multivariate (decoding) analysis in three regions of interest: VC, HPC, vmPFC. Binary decoding was performed for each feature (e.g., colour: red vs green), either by using trials from blocks labelled as Feature RL or Abstract RL. Colour bars represent the mean, error bars the SEM, grey dots individual data points (for each individual, taken as the average across all 3 classifications, i.e., of all feature), * *p*<0.05, ** *p*<0.01, *** *p*<0.001.

The next question we asked was, can we retrieve feature information from HPC and vmPFC activity patterns? In order to abstract and operate in the latent space, an agent is still bound to represent and use the features, because the rules are dictated by features’ combinations. One possibility is that feature information is represented solely in sensory areas. What matters then is the connection with and/or the read out of, vmPFC or HPC. Accordingly, neither HPC or vmPFC should represent feature information (regardless of the strategy used). Alternatively, feature-level information could be represented also in higher cortical regions under Abstract RL to explicitly support (value-based) relational computations. To resolve this question, we applied multivoxel pattern analysis to classify basic feature information (e.g., colour: red vs green) in three regions of interest: the VC, HPC, and vmPFC; separately for trials labelled as Feature RL or Abstract RL. We found the classification accuracy to be significantly higher in Abstract RL than in Feature RL trials in both HPC and vmPFC (two-sided Wilcoxon signed rank test, HPC: z = −4.21, *p*_*(FDR)*_ < 0.001, vmPFC: z = −3.15, *p*_*(FDR)*_ = 0.002, Figure 5C), while there was only a negligible difference in VC (z = −1.30, *p*_*(FDR)*_ = 0.20, Figure 5C). The difference in feature decodability was significantly larger in HPC and vmPFC compared to VC (LMEM model ‘y ~ ROI + (1|participants)’, y: difference in decodability, t_97_ = 2.52, *p* = 0.013). This empirical result supports the second hypothesis: in Abstract RL features are represented in the neural circuitry incorporating the HPC and vmPFC, beyond a simple read out of sensory cortices. In Feature RL, representing feature-level information in sensory cortices alone should suffice because each visual pattern mapped to a task-state.

We expanded on this idea with two searchlight multivoxel pattern analyses. In short, we inquired which brain regions were sensitive to feature relevance (when the feature was relevant to the rule or not), and whether we could recover representations of the latent rule itself (the fruit preference). Besides the occipital cortex, significant reduction in decoding accuracy when a feature was irrelevant to the rule compared to when it was relevant was also detected in the OFC, ACC, vmPFC and dorsolateral PFC (Figure S6A). Multivoxel patterns in the dorsolateral PFC and lateral OFC further predicted fruit class (Figure S6B).

### Artificially injecting value in sensory representations with neurofeedback fosters abstraction

Our computational and neuroimaging results indicate valuation plays a key role in abstraction. Two hypotheses on the underlying mechanism can be outlined here. On one hand, the effect of vmPFC value computations could remain localized within the prefrontal circuitry. For example, this could be achieved by representing and ranking incoming sensory information for further processing within the HPC-OFC circuitry. Alternatively, value computation could determine abstractions by directly affecting early sensory areas - i.e., a top-down (attentional) effect, to ‘tag’ the relevant sensory information (46). Work in rodents has reported strong top-down modulation of sensory cortices by OFC neurons implicated in value computations (47, 48). We thus hypothesized abstraction could result from a direct effect of value in the VC. Therefore, artificially adding value to a neural representation of a task-relevant feature should result in enhanced behavioural abstraction.

Decoded neurofeedback is a form of neural reinforcement based on real time fMRI and multivoxel pattern analysis. It is the closest approximation to a non-invasive causal manipulation, with high specificity and administered without participants’ awareness (49–51). Such reinforcement protocols can reliably lead to behavioural or physiological changes that last over time (52–56). We used this procedure in a follow-up experiment (a subgroup of N = 22 from the main experiment, see Methods) to artificially add value (rewards) to neural representation in VC of a target task-related feature (Figure 6A). At the end of two training sessions, participants completed 16 blocks of the pacman fruit preference task, outside of the scanner. Task blocks could be labelled as ‘relevant’ (8 blocks) if the feature tagged with value was relevant to the block’s rule, or ‘irrelevant’ otherwise (8 blocks).

**Figure 6:**
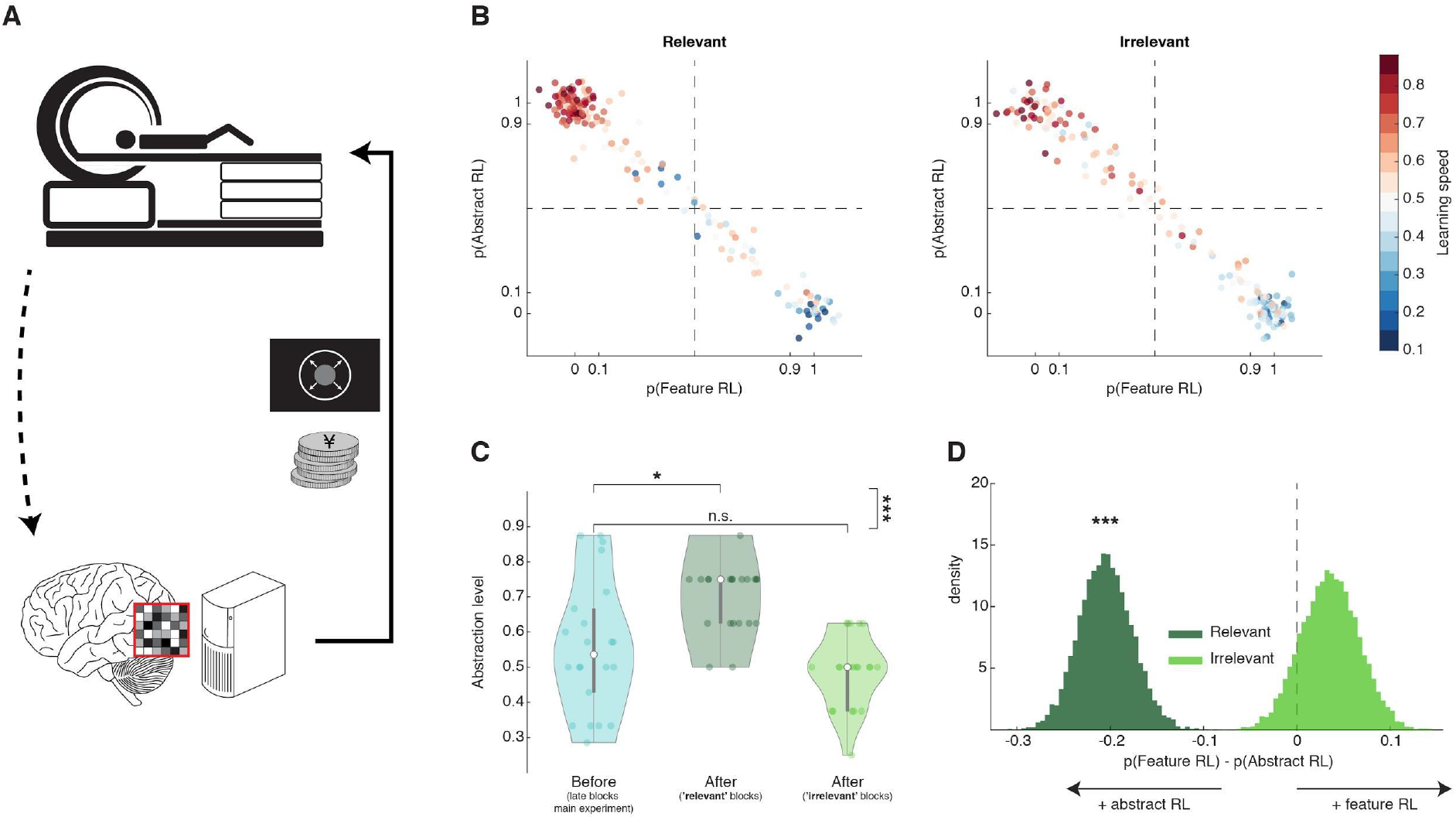
Artificially adding value to a feature’s neural representation. ***A***, Schematic diagram of the follow-up multivoxel neurofeedback experiment. During the neurofeedback procedure, participants were rewarded for increasing the size of a disc on the screen (max session reward 3,000 JPY). Unbeknownst to them, the disc size was changed by the computer program to reflect the likelihood of the target brain activity pattern (corresponding to one of the task features) measured in real time. ***B***, Blocks were subdivided based on the feature targeted by multivoxel neurofeedback: ‘relevant’ or ‘irrelevant’ for the blocks’ rules. The scatter plots replicate the finding from the main experiment, with a strong dependency between Feature RL / Abstract RL and learning speed. Each coloured dot represents a single block, from one participant. Data aggregated from all participants. ***C***, Abstraction level computed for each participant from all blocks belonging to: 1) left, the latter half of the main experiment (as in Figure 3G, but only selecting participants who took part in the multivoxel neurofeedback experiment); 2) centre, post-neurofeedback, for the ‘relevant’ condition; 3) right, post-neurofeedback, for the ‘irrelevant’ condition. Coloured dots represent participants, shaded areas the density plot, central white dots the median, the dark central bar the interquartile range, and thin dark lines the lower and upper adjacent values. ***D***, Bootstrapping the difference between model probabilities on each block, separately for ‘relevant’ and ‘irrelevant’ conditions. * *p*<0.05, *** *p*<0.001

Data from the ‘relevant’ and ‘irrelevant’ blocks were analysed separately. The same model fitting procedure used in the main experiment established whether participants’ choices in the new blocks were driven by a Feature RL or Abstract RL strategy. ‘Relevant’ blocks appeared to be associated with a behavioural shift towards Abstract RL, while there was no substantial qualitative effect in the ‘irrelevant’ blocks (Figure 6B). To quantify this effect, we first applied a binomial test, finding that behaviour in blocks where the targeted feature was relevant displayed markedly increased abstraction (base rate 0.5, number of Abstract RL blocks given total number of blocks; ‘relevant’: P(123|176) = 1.37×10^−7^, ‘irrelevant’: P(90|176) = 0.82). We then measured the abstraction level for each participant and directly compared it to the level attained by the same participants in the late blocks of the main experiments (from Figure 3G). Participants increased their use of abstraction in ‘relevant’ blocks, while no significant difference was detected in the ‘irrelevant’ blocks (Figure 6C, two-sided Wilcoxon signed rank test, ‘relevant’ blocks: z = 2.44, *p* = 0.015, ‘irrelevant’ blocks: z = −1.55, *p* = 0.12, ‘relevant’ vs ‘irrelevant’: z = 4.01, *p* = 6.03×10^−5^). Finally, we measured the difference between model probabilities P_(Feature RL)_ - P_(Abstract RL)_ for each block, and bootstrapped the mean over blocks (with replacement) 10,000 times to generate a distribution for the ‘relevant’ and ‘irrelevant’ conditions. Replicating the results reported above, behaviour in ‘relevant’ blocks had higher probability to be driven by Abstract RL (Figure 6D, perm. test *p* < 0.001), while Feature RL tended to appear more in ‘irrelevant’ blocks. A strategy shift towards abstraction, specific to the blocks in which the target feature was tagged with reward, indicates the effect of value in facilitating abstraction is likely to be mediated by a change in the early processing stage of visual information. In this experiment, by means of neurofeedback, value (in the form of external rewards) ‘primed’ a target feature. Hence, the brain used these ‘artificial’ values when constructing abstract representations by tagging certain sensory channels. Critically, this manipulation indicates that value tagging of early representation has a causal effect on abstraction and consequently on the learning strategy.

## Discussion

The ability to generate abstractions from simple sensory information has been suggested as key to support flexible and adaptive behaviours (4, 10, 11). Here, using computational modelling based on a mixture-of-experts RL architecture, we found that value predictions drive participants’ selection of the appropriate representation to solve the task. Participants explored and used the task dimensionality through learning, as they shifted from a simple feature-based strategy to using more sophisticated abstractions. The more participants used Abstract RL, the faster they were at solving the task. Note that in this task structure learning speed and abstraction are linked: to learn faster, an agent must use Abstract RL – as any other strategy would have resulted in slower completion of the task’s blocks.

These results support the idea that efficient decision-making processes in the brain depend on higher-order, summarized representations of task-states (6). Further, abstraction likely requires a functional link between sensory regions and areas encoding value predictions about task states (Figure 4C, the VC-vmPFC coupling predicted participants’ learning speed). This is in line with previous work that has demonstrated how estimating reward value of individual features provides a reliable and adaptive mechanism in RL (9). We extend this notion by showing that the mechanism may support the formation of abstract representations to be further used for learning computations, for example the selection of the appropriate abstract representation.

There is an important body of work considering how the HPC is involved in the formation and update of conceptual information (14, 24, 26, 57). Likely, the HPC’s role is to store, index and update conceptual/schematic memories (14, 25, 58). The ‘creation’ of new concepts or schemas may happen elsewhere. A good candidate could be the mPFC or vmPFC in humans (58, 59). Our results expose a potential mechanism on how vmPFC interplays with HPC in the construction of goal-relevant abstractions: vmPFC-driven valuation of low-level sensory information serves to channel specific features/components to higher order areas (e.g., the HPC, vmPFC, but also the dorsal prefrontal cortex for instance). In line with this interpretation, we found vmPFC to be more engaged in Abstract RL, while HPC was equally active under both abstract and feature-based strategies (Figure 5B). When a feature was irrelevant to the rule, its decodability from activity patterns in OFC/DLPFC decreased (Figure S6A). These findings dovetail well with the regions’ role in constructing goal-dependent task states and abstract rules from relevant sensory information (28, 60, 61). Furthermore, we found the blocks’ rules to be also encoded in multivoxel activity patterns within the OFC/DLPFC circuitry (Figure S6B) (60, 62, 63).

How these representations are actually used remains an open question. This study nevertheless suggests that there is no single region in the brain that maintains a fixed task state. Rather, the configuration of elements that determines a state is continuously reconstructed over time. At first glance this may appear dispendious and inefficient. But this approach would provide high flexibility in noisy and stochastic environments, and where temporal dependencies exist (as in most real-world situations). By continuously recomputing task states, the agent can make more robust decisions because these are related to the subset of most relevant, up-to-date information. Such a computational coding scheme shares strong analogies with HPC representations, whereby neurons continuously generate representations of alternative future trajectories (64).

One significant aspect of discussion concerns the elements used to construct abstractions. We leveraged simple visual features (colour, or stripe orientation), rather than more complex stimuli or objects that can be linked together conceptually (24, 65). Abstractions happen at several levels, from features, to exemplars, concepts/categories, all the way to rules and symbolic representations. In this work we effectively studied one of the lowest levels of abstraction. One may wonder whether the mechanism we identified here generalizes to more complex scenarios. While our work cannot decisively support this, we are inclined to believe it is unlikely that the brain uses an entirely different strategy to generate new representations at different levels of abstractions. Rather, the internal source of information to be abstracted from should be different, but the algorithm itself should be the same or, at the very least, very similar. The fact our PPI analysis showed a link between vmPFC and VC may point to this distinguishing characteristic of our study: learning through abstraction of simple visual features should be related to early VC. Features in other modalities - e.g., auditory - would involve the functional coupling between auditory cortex and the vmPFC. When learning about more complex objects or categories, we expect to see a stronger reliance on the HPC (14, 24). Future studies could incorporate different levels of complexity, or different modalities, within a similar design so as to directly test this prediction and dissect the exact neural contributions. Depending on which type of information is relevant at any point in time, we suspect different areas will be coupled with the vmPFC to generate value representations.

In our second experiment, we implemented a direct assay to test our (causal) hypothesis that value about features is guiding abstraction in learning. Artificially adding value in the form of reward to a feature representation in the VC resulted in increased use of abstractions. The facilitating effect of value on abstraction can be thus directly linked to changes in the early processing stage of visual information. In line with this interpretation, recent work in mice has elegantly reported how value governs a functional remapping in the sensory cortex by direct lateral OFC projections carrying RPE information (47), as well as by modulating the gain of neurons to irrelevant stimuli (48). While these considerations clearly point to a central role of the vmPFC and valuation in abstraction - by controlling sensory representations, it remains to be investigated whether this effect results in more efficient *construction* of abstract representations, or in better *selection* of internal abstract models.

Given the complex nature of our design, there are some limitations to this work. For example, there isn’t an applicable feature decoder to test actual feature representations (e.g., colour vs orientation) - or the likelihood of one feature against another. In our task design, on every trial, all features were used to define the pacman characters. Furthermore, we did not have a localizer session in which the features were presented in isolation (see Supplementary note 2 for further discussion). Future work could investigate how separate feature representations may emerge on the path to abstractions, for example in the vmPFC, and their relation to feature levels (e.g., for colour: red vs green) as reported here. We speculate that parallel circuits linking the prefrontal cortex and basal ganglia could keep track of these levels of information or abstraction, possibly in a hierarchical fashion (11, 66, 67).

One may point out that what we call ‘Abstract RL’ is, in fact, just an attention-mediated enhanced processing. Yet, if top-down attention were the sole underlying driver in Abstract RL, we contend that the pattern of results would have been different. For example, we would expect to see a marked increase in feature decodability in VC (68, 69). This was not the case here, with only a minimal, non-significant, increase. More importantly, the results of the decoded neurofeedback manipulation question this interpretation. Because decoded neurofeedback operates unconsciously (49, 51), value was added directly at the sensory representation level (limited to the targeted region), precluding alternative top-down effects. That is not to say that attention does not play a significant role in mediating this type of abstract learning; however, we argue that attention is most likely to be an effector of the abstraction and valuation processes (70). A simpler top-down attentional effect was indeed evident in the supplementary analysis comparing feature decoding in the ‘relevant’ and ‘irrelevant’ cases (Figure S6A): occipital regions displayed large effect sizes (irrespective of the learning strategy used to solve the task).

While valuation and abstraction appear tightly associated in reducing the dimensionality of the task space, what is the underlying mechanism? The degree of neural compression in the vmPFC has been shown to relate to features most predictive of positive outcomes, under a given goal (59). An attractive view is that valuation may be interpreted as an abstraction in itself; value could provide a simple and efficient way for the brain to operate on a dimensionless axis. Each point on this axis could be indexing a certain task state, or even behavioural strategy, as a function of its assigned abstract value. The neuronal encoding of value about certain features, or choice options, may help the system construct useful representations that can, in turn, inform flexible behavioural strategies (6, 28, 31).

In summary, this work provided evidence for a role played by valuation that goes beyond the classic view in decision-making and neuroeconomics. We show that valuation subserves a critical function in constructing abstractions. One may go further and speculate that valuation, by generating a common currency across perceptually different stimuli, may be an abstraction in itself, and that it is tightly linked to the concept of task states in decision-making. We believe this work not only offers a new perspective on the role of valuation in generating abstract thoughts, but also reconciles (apparently disconnected) findings in decision-making and memory literature on the role of the vmPFC. In this context value is not a simple proxy of a numerical reward signal but is better thought of as a conceptual representation or schema built on-the-fly to respond to a specific behavioural demand. We believe our findings thus provide a fresh vision on the invariable presence of value signals in the brain that play an important algorithmic role in the development of sophisticated learning strategies.

## END NOTES

## Acknowledgements

We thank Kaori Nakamura, Yasuo Shimada for experimental assistance.

## Authors’ contributions

A.C., P.S., M.K. and B.D.M. contributed to the conceptual development of the work and designed the experiments; P.S. developed and piloted the task under guidance of A.C. and B.D.M.; A.C. and A.Y. collected the data; A.C., A.Y., and M.H. analysed the data; A.C., B.D.M. and M.K. drafted the manuscript; all authors revised the manuscript.

## Materials and correspondence

Please contact A.C. or B.D.M. for correspondence or requests related to materials.

## Methods

### Participants

Forty-six participants with normal or corrected-to-normal vision were recruited for the main experiment (learning task). Based on pilot data and the available scanning time in one session (60 minutes), we set the following conditions of exclusion: failure to learn the association in 3 blocks or more (i.e., reaching a block’s limit of 80 trials without having learned the association), or failure to complete at least 11 blocks in the allocated time. Eleven participants were removed based on the above predetermined conditions. Additionally, 2 more participants were removed due to head motion artifacts. Thus, 33 participants (22.4 ± 0.3 y.o.; 8 females) were included in the main analyses. Of these, 22 participants (22.2 ± 0.3 y.o.; 4 females) returned for the follow-up experiment, based on their individual availability. All results presented up to Figure 5 are from the 33 participants who completed the learning task. All results pertaining to the neurofeedback manipulation are from the subset of 22 participants that were called back.

All experiments and data analyses were conducted at the Advanced Telecommunications Research Institute International (ATR). The study was approved by the Institutional Review Board of ATR. All participants gave written informed consent.

### Learning task

The task consisted of learning the fruit preference of pacman-like characters. These characters were made of 3 different features, each with two levels (colour: green vs red, stripes orientation: horizontal vs vertical, mouth direction: left vs right). On each trial, a character composed of a unique combination of the three features was presented. The experimental session was divided into blocks, on each of which a specific rule directed the association between features and preferred fruits. There were always 2 relevant features and 1 irrelevant feature, but these changed randomly at the beginning of each block. Blocks could thus be of three types: CO (colour-orientation), CD (colour-direction), and OD (orientation-direction). Furthermore, to avoid a simple logical deduction of the rule after 1 trial, we introduced the following pairings. The 4 possible combinations of 2 relevant features with 2 levels were paired with the 2 fruits in both a symmetric or asymmetric fashion - 2×2 or 3×1. The appearance of the 2 fruits was also randomly changed at the beginning of each new block (see Figure 1B, e.g., green-vertical: fruit 1, green-horizontal: fruit 2, red-vertical: fruit 1, red-horizontal: fruit 2 *or* green-vertical: fruit 2, green-horizontal: fruit 2, red-vertical: fruit 1, red-horizontal: fruit 2).

Each trial started with a black screen for 1 sec, following which the character was presented for 2 sec. Then, while the character continued to be present at the centre of the screen, the 2 fruit options appeared below, to the right and left sides (see Figure 1). Participants had 2 sec to indicate the preferred fruit by pressing a button (right for the fruit to the right, left vice versa). Upon registering a participant’s choice, a coloured square appeared around the chosen fruit: green if the choice was correct, red otherwise. The square remained for 1 sec, following which the trial ended with a variable ITI - bringing the trial to a fixed 8 sec duration.

Participants were simply instructed that they had to learn the correct rule on each block, and the rule itself (relevant features + association type) was hidden. Each block contained up to 80 trials, but a block could end before if participants learned the target rule. Learning was measured as a strike of correct trials (between 8 and 12, determined randomly on each block). Participants were instructed that each correct choice added one point, while incorrect choices did not alter the balance. But importantly, they were told that the points obtained would be weighted by the speed of learning on that block. The faster the learning, the greater the net worth of the points - the monetary reward was computed at the end of each block according to the formula:

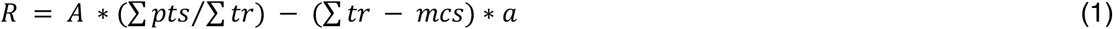

Where *R* is the reward obtained on that block, *A* the maximum available reward (150JPY), Σ*pts* the sum of correct responses, Σ*tr* the number of trials, *mcs* the maximum length of a correct strike (12 trials), and *a* is a scaling factor (*a* = 1.5). This formula ensures a time-dependent decay of the reward that approximately follows a quadratic fit. In case participants completed a block in less than 12 trials, if the amount was larger than 150JPY it was rounded to 150JPY.

The maximum terminal monetary reward over the whole session was set at 3,000 JPY; participants on average earned 1251 ± 46 JPY (blocks in which participants failed to learn the association within the 80 trials limit were not rewarded). For each experimental session there was a sequence of 20 blocks that was pre-generated pseudo-randomly, and on average participants completed 13.6 ± 0.3 blocks. In the post-neurofeedback behavioural test all participants completed 16 blocks, 8 of which had the targeted feature as relevant, while in the other half the targeted feature was irrelevant. The order was arranged pseudo-randomly such that, on the both halves of the session there were 4 blocks of each type. In the post-neurofeedback behavioural session, all blocks only had asymmetric pairings with preferred fruits.

For the sessions done in the MR scanner, participants were instructed to use their dominant hand to press buttons on a dual-button response pad to provide their choices. Concordance between responses and buttons was indicated on the display and, importantly, randomly changed across trials to avoid motor preparation confounds (i.e. associating a given preference choice with a specific button press).

The task was coded with PsychoPy v1.82.01 (71).

### Computational modelling part 1: mixture-of-experts RL model

We built on a standard RL model (1) and prior work in machine learning and robotics to derive the mixture-of-experts architecture (33, 34, 37). In this work the mixture-of-experts architecture is composed of several ‘expert’ RL modules, each tracking a different representational space, and each with its own value function. On each trial, the action selected by the mixture-of-experts RL model is given by the weighted sum of the actions computed by the experts. The weight reflects the responsibility of each expert, which is computed from the SoftMax of the squared prediction error. In this section we define the general mixture-of-expert RL model, and in the next section we define each specific expert, which are based on the different task-state representations being used.

Formally, the mixture-of-expert RL model global action is defined as:

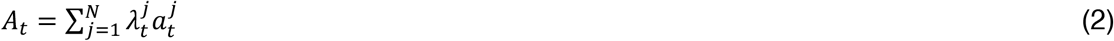

Where *N* is the number of experts, **λ** the responsibility signal, and *a* the action selected by the *j*th-model. Thus, **λ** is defined as:

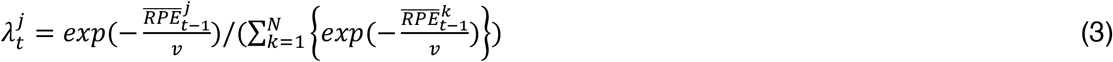

Where *N* is the same as above, ***v*** is the RPE variance. The expert’s uncertainty 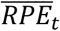 is defined as:

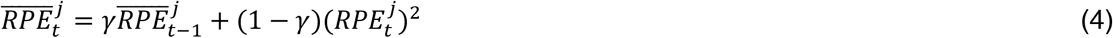

Where **γ** is the forgetting factor, which controls the strength of the impact of prior experience on the current uncertainty estimate. The most recent RPE is computed as:

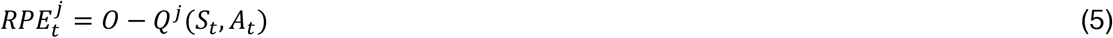

Where *O* is the outcome (reward: 1, no reward: 0), *Q* is the value function, *S* the state for the current expert, and *A* the global action computed with equation (2). The update to the value function can therefore be computed as:

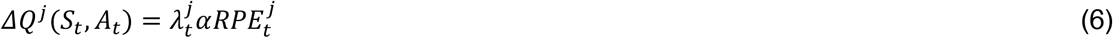

Where **λ** is the responsibility signal computed with equation (3), **α** is the learning rate (assumed equal for all experts), and *RPE* computed with equation (5). Finally, for each expert, the action *a* at trial *t* is taken as the argmax of the value function as follows:

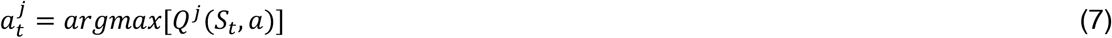

Where *Q* is the value function, *S* the state at current trial, and *a* the two possible actions.

### Computational modelling part 2: Feature RL and Abstract RLs

Each (expert) RL algorithm is built on a standard RL model (1) to derive variants that differ on the number and type of states visited. Here, a state is defined as a combination of features. Feature RL has 2^3^ = 8 states, where each state was given by the combination of all three features (e.g. colour, stripe orientation, mouth direction: green, vertical, left). Abstract RL is designed with 2^2^ = 4 states, where each state was given by the combination of two features.

Note that the number of states does not change for different blocks, only the features used to determine them. These learning models create individual estimates of how participants’ action-selection was dependent on the features they attended and their past reward history. Both RL models share the same underlying structure and are formally described as:

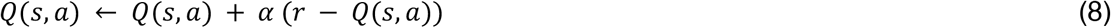

Where *Q*(*s*, *a*) in (8) is the value function of selecting either fruit-option *a* for packman-state *s*. The value of the action selected on the current trial is updated based on the difference between the expected value and the actual outcome (reward or no reward). This difference is called the reward prediction error (RPE). The degree to which this update affects the expected value depends on the learning rate *α*. The larger *α*, the more recent outcomes will have a strong impact. On the contrary, for small *α* recent outcomes will have little effect on expectations. Only the value function of the selected action - which is state-contingent in (8) - is updated. The expected values of the two actions are combined to compute the probability *p* of predicting each outcome using a SoftMax (logistic) choice rule:

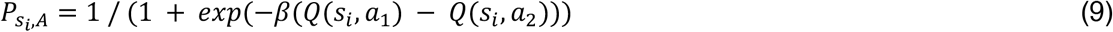

The greediness hyperparameter *β* controls how much the difference between the two expected values for *a_1_* and *a_2_* actually influence choices.

### Procedures for model fitting, simulations and hyperparameter recovery

Hierarchical Bayesian Inference (HBI) was used to fit the models to participants’ behavioural data, enabling precise estimate of the hyperparameters at the block level for each participant (39). The hyperparameters were selected by maximizing the likelihood of the estimated actions given the true actions. For the mixture-of-experts architecture, we fit the model on all participants block-by-block to estimate the hyperparameters at the single block and single participant level. For the subsequent direct comparison between Feature RL and Abstract RL models, we used HBI for concurrent model fitting and comparison on an individual, block-by-block basis. The model comparison provided the likelihood that each RL algorithm best explained participants’ choice data. That is, this measure was a proxy to whether learning followed a Feature RL or Abstract RL strategy. Therefore, for all fitting analyses, blocks were first sorted according to their length, from longer to shorter, at the individual participant’s level. The HBI procedure was then implemented on all participants’ data, proceeding block-by-block.

We also simulated the models’ action-selection behaviour to benchmark their similarity to human behaviour and, in the case of the Feature RL vs Abstract RL comparisons, to additionally compare their formal learning efficiency. In the case of the mixture-of-experts RL architecture, we simply used the estimated hyperparameters to simulate 45 artificial agents, each completing 100 blocks. The simulation allowed us to compute - for each expert RL module - the mean responsibility signal, and the mean expected value across all states for the chosen action. Additionally, we also computed the learning speed (time to learn the rule of a block) and compare it with the learning speed of human participants.

In the case of the simple Feature RL and Abstract RL models, we added noise to the state information in order to have a more realistic behaviour (from the perspective of human participants). In the empirical data, the action (fruit selection) in the first trial of a new block was always chosen at random because participants did not have access to which were the appropriate representations (states). In later trials participants may have followed specific strategies. For the model simulations we simply assumed that states were corrupted by a decaying noise function:

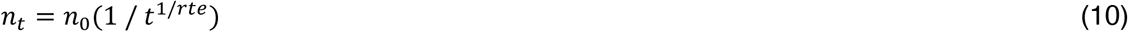

where *n*_*t*_ is the noise level at trial *t*, *n*_*0*_ the initial noise level (randomly drawn from a uniform distribution within the interval [0.3 0.7]), and *rte* was the decay rate, which was set to 3. This meant that in early trials in a block there was a higher chance of basing the action on the wrong representation (at random), rather than following the appropriate value function. The actions in later trials had a decreasing probability of being chosen at random. This approach is a combination of the classic ***ɛ***-greedy policy and standard SoftMax action-selection policy in RL. The hyperparameters values were sampled from the set obtained participants’ data maximum likelihood fits. We simulated 45 artificial agents solving 20 blocks each. The procedure was repeated 100 times for each block with new random seeds. We used two metrics to compare the efficiency of the two models: learning speed (same as above, the time to learn the rule of a block), as well as the fraction of failed blocks (blocks in which the rule was not learned with the 80-trials limit).

We performed a parameter recovery analysis for the simple Feature RL and Abstract RL models based on the data from the main experiment. The parameter recovery analysis was done in order to confirm that the fitted hyperparameters had sensible values and the models themselves were a sensitive description of human choice behaviour (41). Using the same noisy procedure described above, we generated one more simulated dataset, using the exact blocks that were presented to the 33 participants. The blocks from simulated data were then sorted according to their length, and the hyperparameters α and β were fitted by maximizing the likelihood of the estimated actions given the true model actions. We report in Figure S3 the scatter plot and correlation between hyperparameters estimated from participants data and recovered hyperparameters values, showing good agreement notwithstanding the noise in the estimation.

For the data from the behavioural session after multivoxel neurofeedback, blocks were first divided into whether the targeted feature was relevant or irrelevant to the rule of a given block. We then applied the HBI procedure as described above, on all participants, with all blocks of the same type (e.g. targeted feature relevant) ordered by length. This allowed us to compute - based on whether the targeted feature was relevant or irrelevant, the difference in frequency between the models. We resampled with replacement to produce distributions of mean population bias for each block type.

### fMRI scans: acquisition and protocol

All scanning sessions took place in a 3T MRI scanner (Siemens, Prisma) with a 64-channel head coil in the ATR Brain Activation Imaging Centre. Gradient T2*-weighted EPI (echoplanar) functional images with blood-oxygen-level-dependent (BOLD) sensitive contrast and multi-band acceleration factor 6 were acquired. Imaging parameters: 72 contiguous slices (TR = 1 sec, TE = 30 ms, flip angle = 60 deg, voxel size = 2×2×2 mm^3^, 0 mm slice gap) oriented parallel to the AC-PC plane were acquired, covering the entire brain. T1-weighted images (MP-RAGE; 256 slices, TR = 2 s, TE = 26 ms, flip angle = 80 deg, voxel size = 1×1×1 mm^3^, 0 mm slice gap) were also acquired at the end of the first session. For participants who joined the neurofeedback training sessions, the scanner was realigned to their head orientations with the same parameters on all sessions.

### fMRI scans: standard and parametric general linear models

BOLD-signal image analysis was performed with SPM12 (http://www.fil.ion.ucl.ac.uk/spm/), running on MATLAB v9.1.0.96 (r2016b) and v9.5.0.94 (r2018b). The fMRI data for the initial 10 sec of each block were discarded due to possible unsaturated T1 effects. Raw functional images underwent realignment to the first image of each session. Structural images were re-registered to mean EPI images and segmented into grey and white matter. The segmentation parameters were then used to normalize (MNI) and bias-correct the functional images. Normalized images were smoothed using a Gaussian kernel of 7 mm full-width at half-maximum.

GLM1: regressors of interest included ‘High value‘, ‘Low value’ (trials were labelled as such based on the median split of the trial-by-trial expected value for the chosen option computed from the best fitting algorithm - Feature RL or Abstract RL), ‘Feature RL’, ‘Abstract RL’ (trials were labelled as such based on the best fitting algorithm at the block level). For all, we generated boxcar regressors at the beginning of the visual stimulus (character) presentation, with duration 1 sec. Contrast of [1 −1] or [−1 1] were applied to the regressors ‘High value’ - ‘Low value’, and ‘Feature RL’ - ‘Abstract RL’. Specific regressors of no interest included the time in the experiment: ‘early’ (all trials falling within the first half of the experiment), and ‘late’ (all trials falling in the second half of the experiment). The early/late split was done according to the total number of trials: taking as ‘early’ the trials from the first block onward, adding blocks until the trial sum exceeded the total trials number divided by two.

GLM2 (PPI): the seed was defined as a sphere (radius = 6mm) centred around the individual peak voxel from the ‘High value’ > ‘Low value’ group-level contrast, within the vmPFC (peak coordinates [2 50 −10], radius 25 mm). The ROI mask was defined individually to account for possible differences across participants. Two participants were excluded from this analysis as they did not show any significant cluster of voxels in the bounding sphere (even at very lenient thresholds). The GLM for the PPI included 3 regressors (the PPI, the mean BOLD signal of the seed region, and the psychological interaction), as well as nuisance regressors described below.

For all GLM analyses, additional regressors of no interest included a parametric regressor for reaction time, regressors for each trial event (fixation, fruit options presentation, choice, button press, ITI), block regressors, the six head motion realignment parameters, framewise displacement (FD) computed as the sum of the absolute values of the derivatives of the six realignment parameters, the TR-by-TR mean signal in white matter and TR-by-TR mean signal in cerebrospinal fluid.

Second-level group contrasts from all models were calculated as one-sample *t*-tests against zero for each first-level linear contrast. Statistical maps were z-transformed, and then reported at a threshold level of *P(fpr)* < 0.001 (*Z* > 3.09, false positive control meaning of cluster forming threshold), unless otherwise specified. Statistical maps were projected onto a canonical MNI template with MRIcroGL [https://www.nitrc.org/projects/mricrogl/] or a glassbrain MNI template with Nibabel 2.5.0, part of the NiPy suite.

### fMRI scans: pre-processing for decoding

As above, the fMRI data for the initial 10 sec of each run were discarded due to possible unsaturated T1 effects. BOLD signals in native space were pre-processed in MATLAB v7.13 (R2011b) (MathWorks) with the mrVista software package for MATLAB [http://vistalab.stanford.edu/software/]. All functional images underwent 3D motion correction. No spatial or temporal smoothing was applied. Rigid-body transformations were performed to align the functional images to the structural image for each participant. One region of interest (ROI), the HPC, were anatomically defined through cortical reconstruction and volumetric segmentation using the Freesurfer software [http://surfer.nmr.mgh.harvard.edu/]. Furthermore, VC subregions V1, V2, and V3 were also automatically defined based on a probabilistic map atlas 70. The vmPFC ROI was defined based on the significant voxels from the GLM1 ‘High value’ > ‘Low value’ contrast at *p*(fpr) < 0.0001, within the OFC. All subsequent analyses were performed using MATLAB v9.5.0.94 (r2018b). Once ROIs were individually identified, time-courses of BOLD signal intensities were extracted from each voxel in each ROI and shifted by 6 sec to account for the hemodynamic delay (assumed fixed). A linear trend was removed from the time-courses, which were further z-score normalized for each voxel in each block to minimize baseline differences across blocks. The data samples for computing the individual feature identity decoders were created by averaging the BOLD signal intensities of each voxel over 2 volumes, corresponding to the 2 sec from stimulus (character) onset to fruit options onset.

### Decoding: multivoxel pattern analysis (MVPA)

All MVP analyses followed the same procedure. We used sparse logistic regression (SLR) (73), to automatically select the most relevant voxels for the classification problem. This allowed us to construct several binary classifiers (e.g. feature id.: colour - red vs green, stripes orientation - horizontal vs vertical, mouth direction - right vs left).

Cross-validation was used for each MVPA by repeatedly subdividing the dataset into a “training set” and a “test set” in order to evaluate the predictive power of the trained (fitted) model. The number of folds was set at k=50. On each fold, 80% of the data was assigned to the training set, and the remaining to the test set. The samples assigned to either set were randomly chosen in each fold. Furthermore, SLR classification was optimized by using an iterative approach (74): in each fold of the cross-validation, the feature-selection process was repeated 10 times. On each iteration, the selected features (voxels) were removed from the pattern vectors, and only features with unassigned weights were used for the next iteration. At the end of the cross-validation, the test accuracies were averaged for each iteration across folds, in order to evaluate the accuracy at each iteration. The number of iterations yielding the highest classification accuracy was then used for the final computation. Results (Figure 5C) report the cross-validated average of the best yielding iteration.

For the multivoxel neurofeedback experiment, we used the entire dataset to train the classifier in VC. Thus, each classifier resulted in a set of weights assigned to the selected voxels; these weights could be used to classify any new data sample - and therefore, compute a likelihood of it belonging to the target class.

### Real-time multivoxel neurofeedback and fMRI pre-processing

As in previous studies (52, 53, 75), during the multivoxel neurofeedback manipulation, participants were instructed to modulate their brain activity, in order to enlarge a feedback disc and maximize their cumulative reward. Brain activity patterns measured through fMRI were used in real time to compute the feedback score. Unbeknownst to participants, the feedback score, ranging from 0 to 100 (empty to full disc), represented the likelihood of a target pattern occurring in their brain at measurement time. Each trial started with an induction period of 6 sec, during which participants viewed a cue (small grey circle) which instructed them to modulate their brain activity. This period was followed by a 6 sec rest interval, and then by a 2 sec feedback, during which the disc appeared on the screen. Lastly, each trial ended with a 6 sec inter-trial interval (ITI). Each block was composed of 12 trials, and one session could last up to 10 blocks (depending on time availability). Participants did 2 sessions on consecutive days. Within a session the maximum monetary bonus was 3,000 JPY.

The feedback was calculated through the following steps. In each block, the initial 10 sec of fMRI data were discarded to avoid unsaturated T1 effects. First, newly measured whole-brain functional images underwent 3D motion correction using Turbo BrainVoyager (Brain Innovation, Maastricht, Netherlands). Second, time-courses of BOLD signal intensities were extracted from each of the voxels identified in the decoder analysis for the target ROI (VC). Third, the time-course was detrended (removal of linear trend), and z-score normalized for each voxel using BOLD signal intensities measured up to the last point. Fourth, the data sample to calculate the target likelihood was created by taking the average BOLD signal intensity of each voxel over the 6 sec (6 TRs) ‘induction’ period. Finally, the likelihood of each feature level (e.g. right vs left mouth direction) being represented in the multivoxel activity pattern was calculated from the data sample using the weights of the previously constructed classifier.

### Data and code availability

Behavioural data, group-level maps of brain activation, and custom code used to generate the results and figures will be made available upon publication at https://github.com/BDMLab.

