## Supplementary information for "Value Shapes Abstraction During Learning"

**Note 1. Control regressors for main GLM.** By including regressors for ‘early’, ‘late’, ‘High value’, ‘Low value’ together with ‘Feature RL’ and ‘Abstract RL’, the GLM explicitly controlled for two idiosyncratic features of the task - the fact that, overall, Abstract RL blocks had higher expected value (Fig. S3B), and that Abstract RL blocks were also more likely to be found in the latter half of the experiment (Fig. 3F-G and Fig. S2). Moreover, although Abstract RL blocks were associated with higher expected value compared with Feature RL blocks (Fig. S3B), at the trial level value (high / low) and learning strategy (Feature RL or Abstract RL) were uncorrelated (Fig. S6A), thus confirming the regressors’ orthogonality.

**Note 2. Levels of multivoxel fMRI neurofeedback.** It is worth noting that the neurofeedback procedure targeted one feature’s level, e.g., red colour, rather than colour overall. One might wonder why this approach would work nevertheless? Given previous work with fMRI-based decoded neurofeedback (1), the main driver of the effect was most likely due to change in processing in VC, leading to increased functional representation of task features also in PFC (particularly, in vmPFC). Because in the current work feature levels were intrinsically coupled in task space, e.g., if red-horizontal corresponded to fruit 1, then green-vertical too, enhanced processing of red should also directly influence the paired colour.

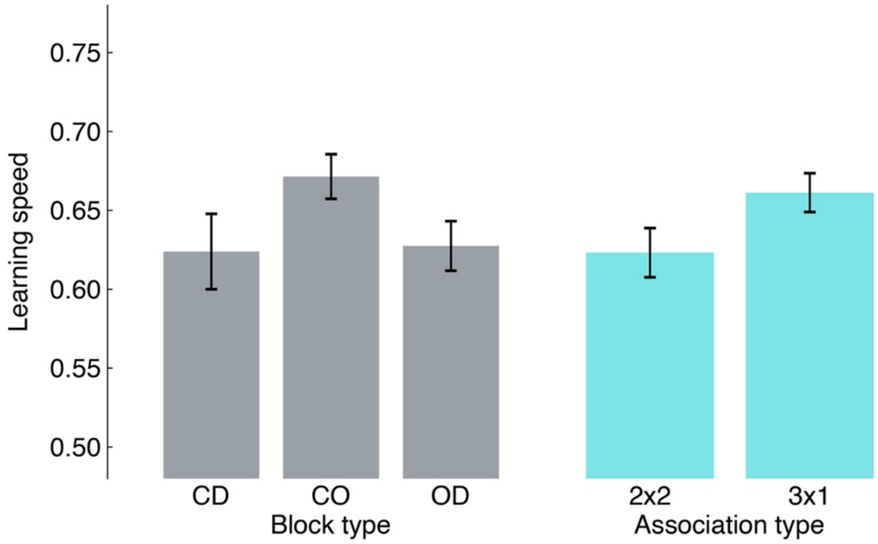


**Figure S1. A, Minimal influence of block type on learning speed.** Average learning speed was computed by pooling block-wise learning speed from all participants for each block or association type. None of the pairwise tests survived multiple comparison correction (FDR). Before correction, the pair CO - OD (block type) had p = 0.047, and the pair 2x2 - 3x1 (association type) had p = 0.065. Bars represent the population mean, error bars the SEM.

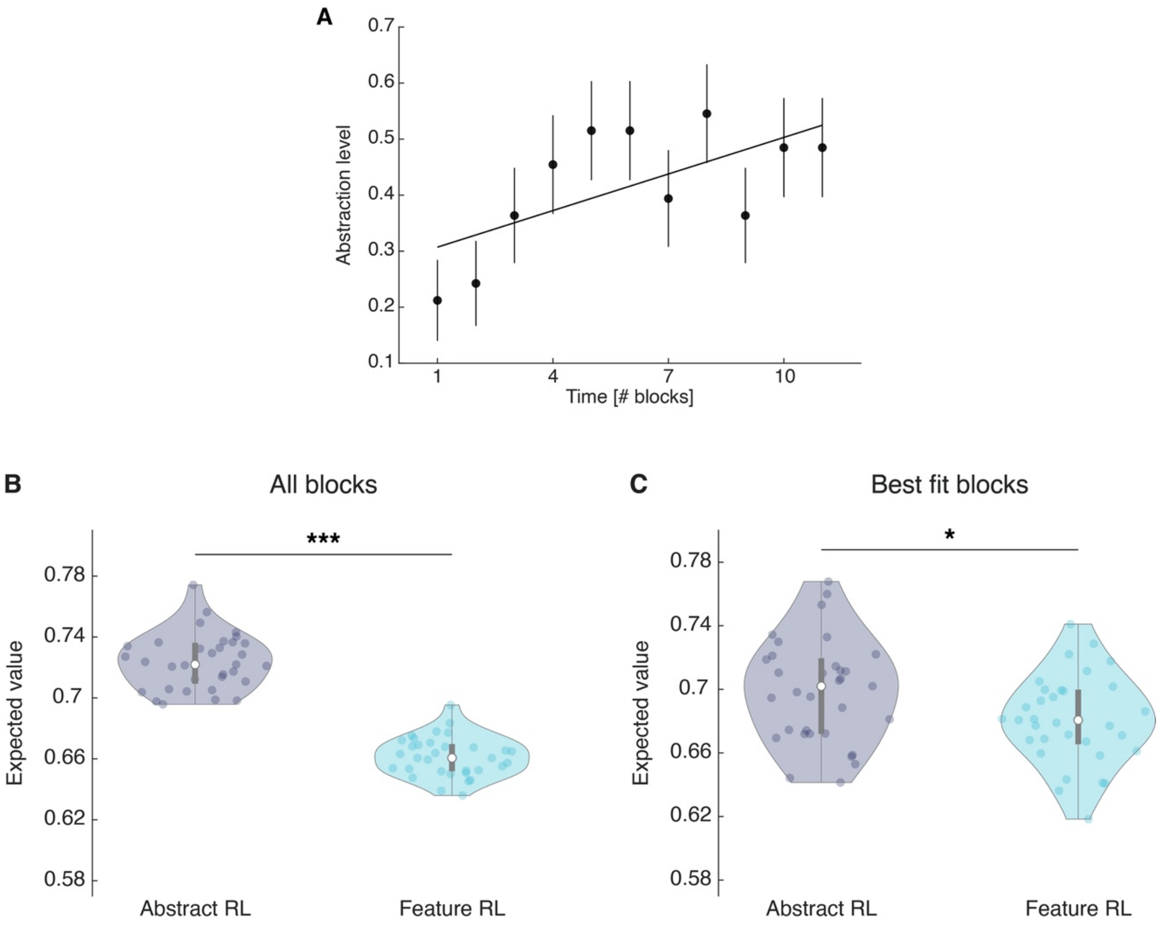


**Figure S2. Abstraction index for single blocks and expected value for the chosen action in Abstract RL and Feature RL. A**, Abstraction index was computed by allocating a value of 0 to blocks labelled as 'Feature RL' and 1 to blocks labelled as 'Abstract RL', for each participant. This was done for the first 11 blocks, which were completed by all participants. The plot indicates that, at the group level, there was a tendency towards increasingly higher abstraction later in time. Each dot represents the population mean, error bars the SEM. The least square line fit, robust regression, and p-value were computed on the 11 mean data points. Robust regression slope = 0.022, t31 = 2.34, P = 0.044.**B**, Mean expected value computed from all blocks, when fitted with either Abstract RL or Feature RL. **C**, Mean expected value computed from the best fitting algorithms on a given block. That is, Abstract RL (resp. Feature RL) refers to the expected value for the chosen action in blocks where Abstract RL (resp. Feature RL) was the best fitting algorithm. One outlier was removed. Each coloured dot represents the average expected value for the chosen options for a single participant - in either Abstract RL (grey) or Feature RL (cyan). Shaded areas represent the density plot, central white dot the median, the dark central bar the interquartile range, and thin dark lines the lower and upper adjacent values. **B**: two-sided t-test, t30 = 35.66, p = 4.03x10-26. **C**: two-sided t-test, t31 = 2.31, p = 0.028.

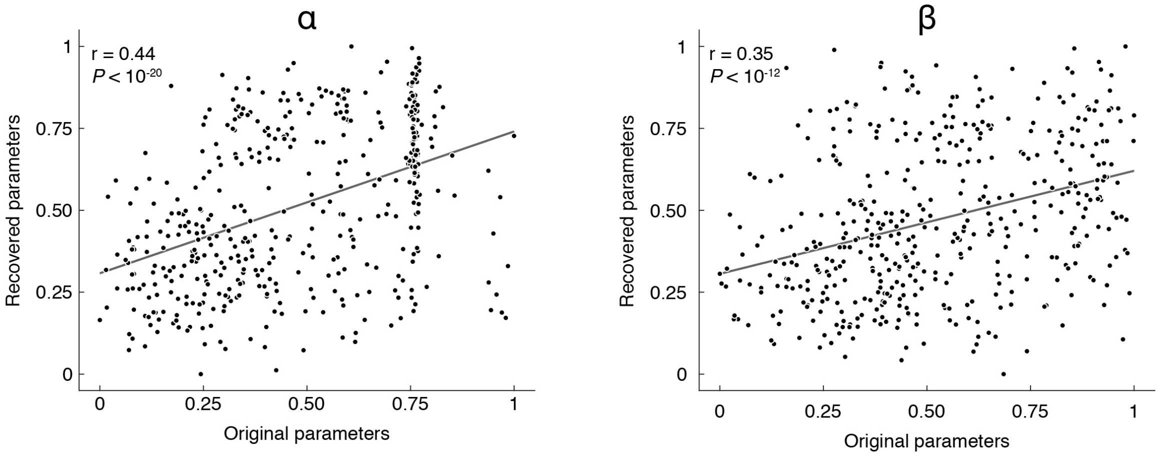


**Figure S3. Parameter recovery.** We first simulated choice data through the models, using the best-fitting parameters. Simulated data was then fed again to the fitting procedure using HBI, separately for each model, in the presence of noise (the update was sometimes not done for the real, correct state but rather for an alternative, random state. Parameters were recovered for each participant, block, and model. Recovered and original parameter values were then pooled across the models and plotted. Values were normalized in the interval [0, 1] for visualization purposes (note that the normalization does not affect the strength of the correlation).

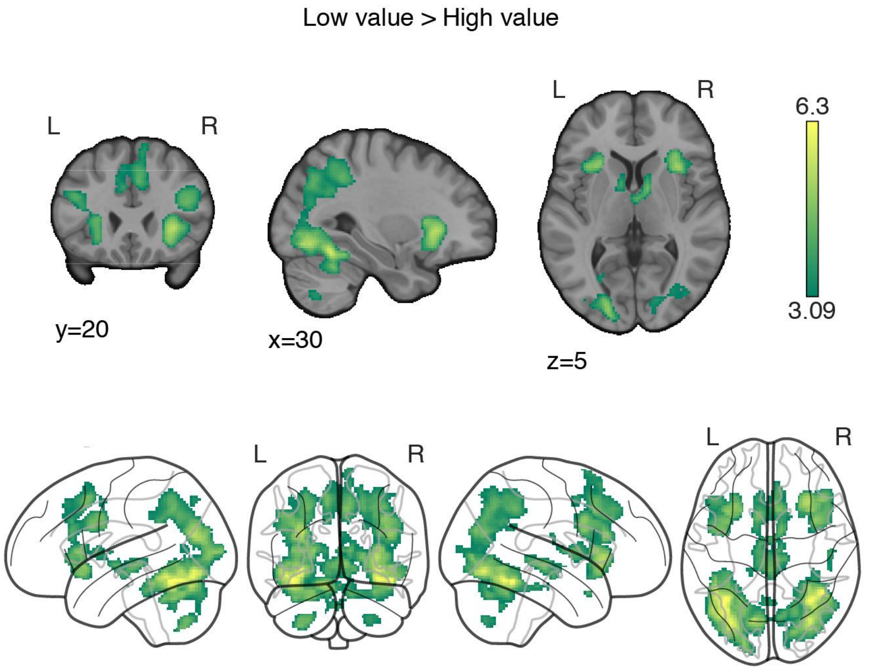


**Figure S4. ‘Low value’ > ‘High value’ GLM contrast.** Neural correlates of (predicted) low value at visual stimulus presentation time. Trials were labelled according to a median split of the expected value for the chosen option as computed by the best fitting model, at the participants and block level. The statistical parametric map was z-transformed, and false-positive means of cluster formation (fpr) correction was applied. p(fpr) < 0.001, Z > 3.09.

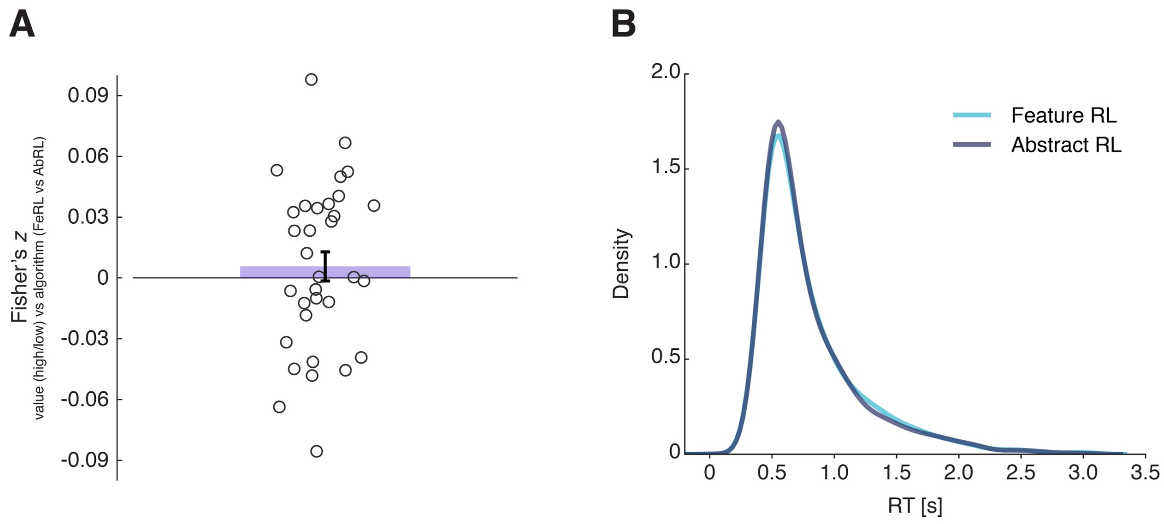


**Figure S5. Reaction time in Feature RL and Abstract RL trials. A**, Fisher-transformed coefficients (Spearman $\rho$) of the correlation between high/low value trials and Feature RL / Abstract RL trials. The coloured bar represents the population mean, the error bar the SEM, and dots individual participants' data. The two labels used in the main GLM appear generally uncorrelated. Wilcoxon sign rank test against median 0, z = 0.72, P = 0.47. **B**, Reaction time (RT) data pooled over all participants for trials in blocks labelled as 'Feature RL' or 'Abstract RL'. RT was not significantly different between the two strategies. Wilcoxon sign rank test between the two distributions, z = 1.48, p = 0.14.

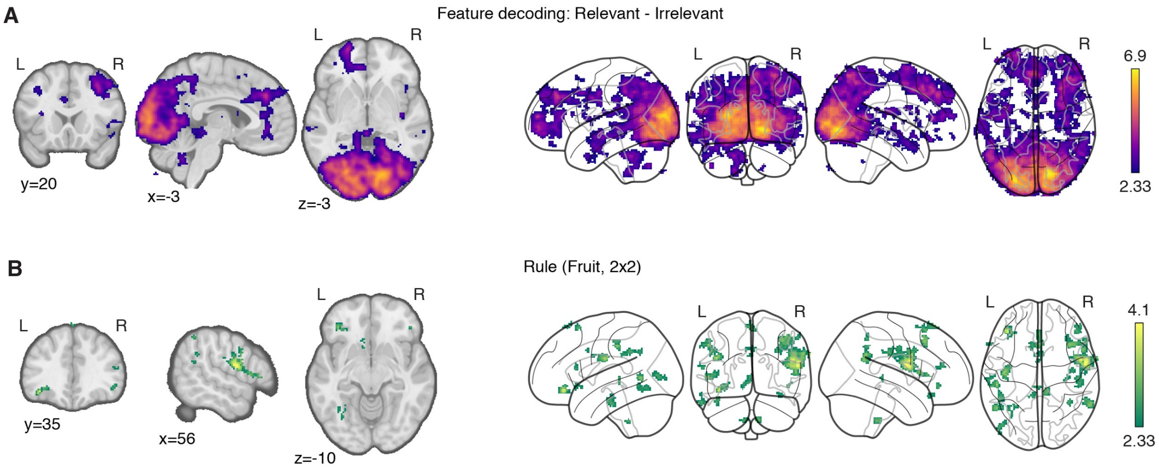


**Figure S6. Multivariate searchlight analyses. A**, Classification was performed for each feature pair (e.g., colour: red vs green), separately for blocks in which the feature in question was relevant or irrelevant to the block's rule. The statistical map reported here represents the strength of the reduction in decoding between trials in which the feature was relevant compared with irrelevant, averaged over all features and participants. **B**, Classification of rule (2x2 only, since 3x1 would have an overlapping feature). For each participant classification was performed as fruit 1 vs fruit 2. All statistical parametric maps were z-transformed, and false-positive means of cluster formation (fpr) correction was applied. $p(fpr) < 0.01$, $Z > 2.33$.
